# Task-specific sensory coding strategies matched to detection and discrimination behaviors in *Apteronotus leptorhynchus*

**DOI:** 10.1101/112995

**Authors:** K.M. Allen, G. Marsat

## Abstract

All sensory systems must reliably translate information about the environment into a neural code, mediating perception. The most relevant aspects of stimuli may change as behavioral context changes, making efficient encoding of information more challenging. Sensory systems must balance rapid detection of a stimulus with perception of fine details that enable discrimination between similar stimuli. We show that in a species of weakly electric fish, *Apteronotus leptorhynchus,* two coding strategies are employed for these separate behavioral tasks. Using communication signals produced in different contexts, we demonstrate a strong correlation between neural coding strategies and behavioral performance on a discrimination task. Extracellular recordings of pyramidal cells within the electrosensory lateral line lobe of alert fish show two distinct response patterns, either burst discharges with little variation between different signals of the same category, or a graded, heterogeneous response that contains enough information to discriminate between signals with slight variations. When faced with a discrimination-based task, the behavioral performance of the fish closely matches predictions based on coding strategy. Comparisons of these results with neural and behavioral responses observed in other model systems suggest that our study highlights a general principle in the way different neural codes are utilized in the sensory system.

**SIGNIFICANCE STATEMENT:** Research relating the structure of stimuli to the response of sensory neurons has left us with a detailed understanding of how different neural codes can represent information. Although various aspects of neural responses have been related to perceptual abilities, general principles relating behavioral tasks to sensory coding strategies are lacking. A major distinction can be made between signals that must simply be detected versus stimuli that must also be finely discriminated and evaluated. We show that these two different perceptual tasks are systematically matched by distinct neural coding strategies and we argue that our study identifies a general principle that is observed in various sensory systems.

**Conflict of interest statement:** The authors declare no competing financial interests.

## INTRODUCTION

Most animals must navigate complex environments, coping with constant sensory input. Being successful requires being adequately prepared to cope with unpredictable stimuli of varying intensity, structure, and behavioral salience. Sometimes an organism merely needs to detect a specific signal within a continuous and noisy sensory stream (1–5). Alternatively, the organism might need to finely evaluate the properties of the signals in order to discriminate meaningful variations(3–5). Our goal is to demonstrate that these two different sensory needs – detection vs. discrimination- are met with different, uniquely optimized, sensory coding schemes.

Deciphering the relationship between the activity of sensory neurons and perception can be divided into three challenges: 1-understanding how relevant sensory information is represented in the neural code; 2-probing sensory abilities through behavior; and 3- correlating the two into a cohesive understanding of “perception”. The relationship between the sensory codes and sensory abilities of an organism has been explored in several systems (7–10). In the visual and vibrissae systems neural codes change as adaptation to a stimulus sets in. After adaptation the initial low-threshold detectability of a novel stimulus is traded for increased discriminability between similar stimuli (11, 12) a change reflected in the behavioral performance of the animal (13). Two neural coding schemes have repeatedly been shown to provide streams of information for the detection vs. discrimination of signals. The first relies on synchronous high frequency firing – typically bursting– over a population, for detection of important stimulus features (14, 15). The second relies on graded responses with heterogeneous firing across the population to support the evaluation and discrimination of fine features of the stimulus (16–18). The relationship between these two strategies and stimulus encoding is firmly established, but we have yet to demonstrate that they indeed are systematically mediating different behavioral tasks. Here we use the weakly electric fish communication system to demonstrate a close association between the stimulus encoding method and the associated behavior. We propose a general principle linking differing sensory codes to two different perceptual tasks: detection vs. discrimination.

Weakly electric fish communication signals, chirps, are transient modulations of their ongoing oscillating electric field, or EOD (electric organ discharge). Two fish interacting will perceive each other’s field as quasi-sinusoidal modulations of their own EOD –called beats– and chirps as transient disruptions of this regular background. In typical male-female courtship interactions, signals will consist of high frequency beats with males producing big (type 1) chirps (19). Male-male aggressive interactions elicit the production of small (type 2) chirps that typically interrupt a low-frequency beat (20). Neurophysiological recordings of the primary electrosensory area in the brain, the electrosensory lateral line lobe (ELL), show that small chirps on low frequency beats are encoded by synchronized bursting among the population of pyramidal cells consistent with a feature detection code. Courtship signals, big chirps on high frequency beats, are encoded via graded, heterogeneous firing (21). This latter response type, but not the former, is well suited to support the efficient discrimination of small variations in chirp properties.

Although the signals described above are typical of courtship and aggression respectively, signaling patterns are presumably more diverse than previously reported. Recent field studies report the use of small chirps paired with high frequency beats during courtship (22), behavior not previously seen in lab studies. Thus, this system offers a unique opportunity to ask the following question: is the neural code employed simply a function of the signal or is it consistently matched to the behavioral task in order to enhance either the capacity to discriminate or simply detect the stimulus?

## RESULTS

In this study, three types of communication signals were used: low frequency beats (10 Hz, typical of agonistic encounters) paired with small chirps and high frequency beats (120 Hz, typical of courtship) with either big chirps or small chirps. We presented several chirp variants, differing in duration and in frequency rise within a range typical of *A. leptorhynchus* signals for each chirp type. We used a spike metrics distance and ROC analysis to characterize the similarity in response pattern between responses of a population of pyramidal neurons (up to 20 ON or OFF cells) to a given chirp stimulus and across responses to variants (Fig S1). The result is presented as percent error (Figures 1 and 2) representing how often an ideal observer would classify a chirp incorrectly as a function of the number of neurons included in the analysis. Efficient discrimination should be accurate even when based on only a few neurons. Alternatively, the pooled responses from many neurons may not contain enough information to accurately discriminate between two stimuli.

**Figure 1:**
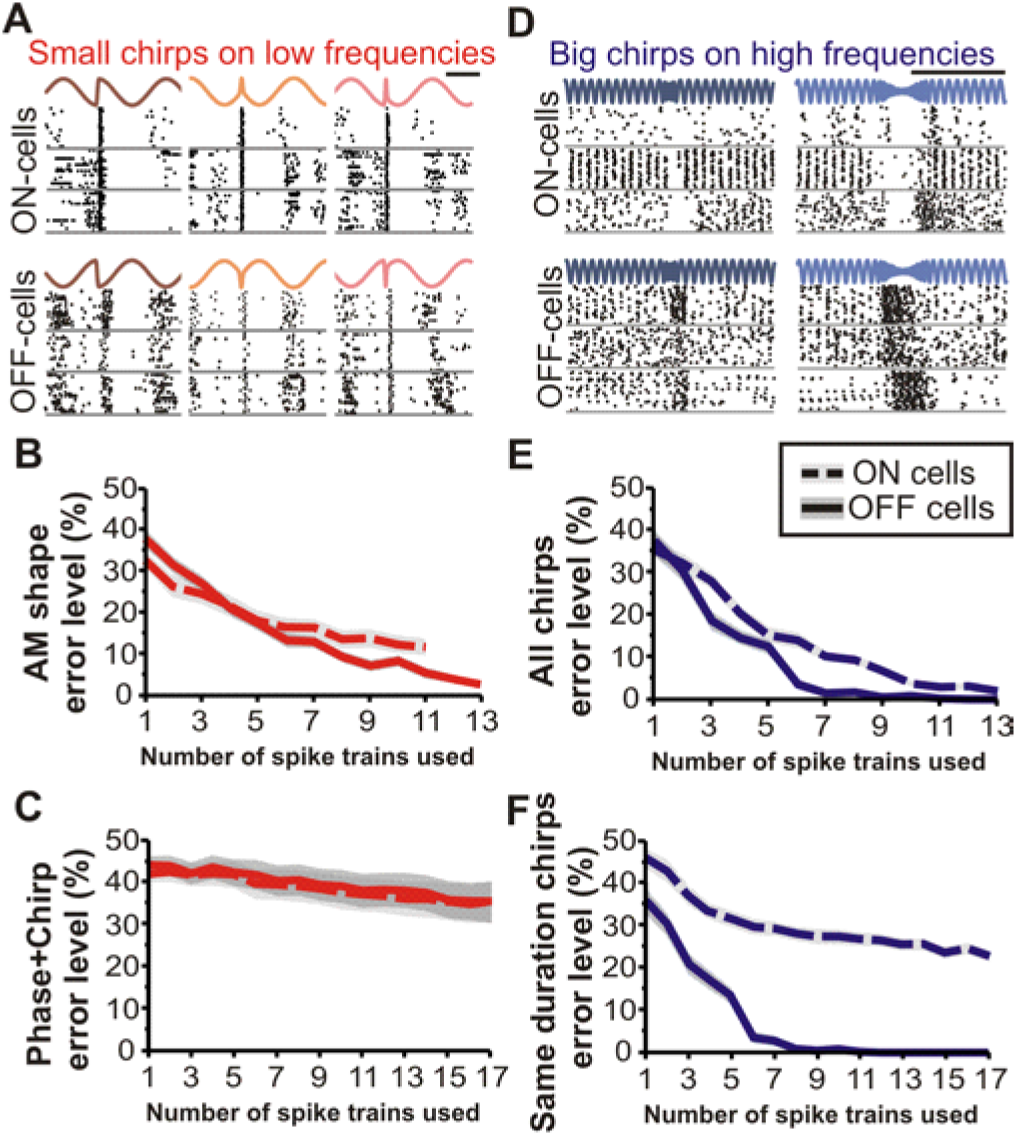
Small chirps cannot be discriminated when phase is varied on low frequency beats; big chirps on high frequency beats can be discriminated. (A) Example responses of four ON-cells presented with small chirps on a 10Hz beat. The phase at which the chirp occurs changes the shape of the signal, even when the chirp frequency and duration are the same. The first and the third chirp stimuli have identical chirps (same frequency and duration) but occurred at different phases; the second chirp differs in frequency and duration from the other two but occurred at the same phase as the third chirp in this panel. (B) Discrimination analysis of chirp identity performed on spike train responses to chirps varying in both duration and frequency but occurring at the same phase. The shape of the AM stimulus is discriminated better by OFF-cells than ON-cells. Shaded area indicates standard error. (C) For chirps with varying parameters and phases, as in natural interactions, error level remains high. (D) Example responses of both ON- and OFF-cells to big chirps that vary in both frequency and duration on a 120Hz beat. Big chirps variably increase OFF-cell firing rate; ON-cells are inhibited by big chirps. (E) Chirps that vary in both duration and frequency rise are discriminated by both ON- and OFF-cells. (F) Chirps that vary only in frequency, but not duration, can only be discriminated by the graded responses of OFF-cells at the number of spike trains examined.

**Figure 2:**
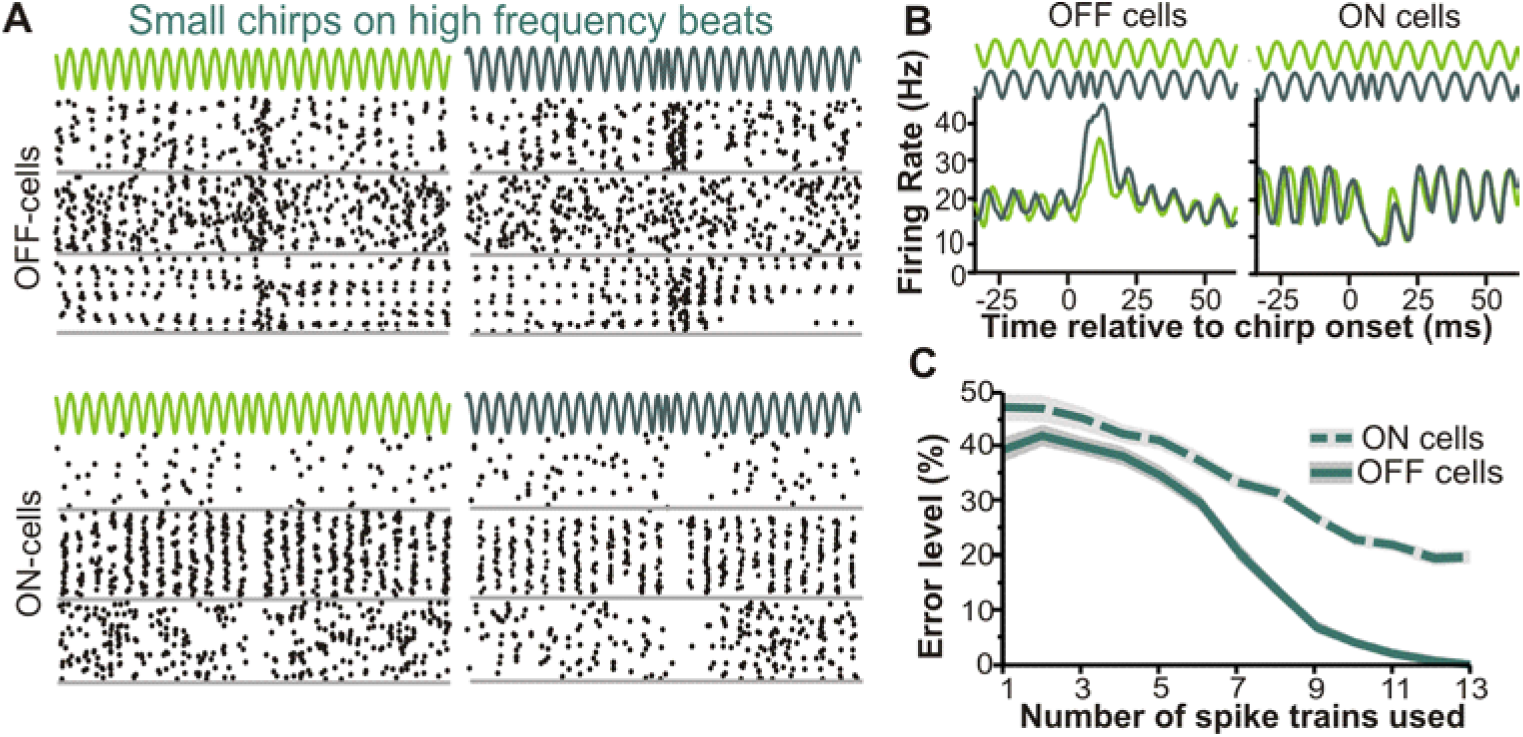
Small chirps on high frequency beats can be discriminated. (A) Example responses of OFF-(top) and ON-cells (bottom) to small chirps on 120Hz beats. Similar to big chirps, OFF-cells show varied increases in firing rate, and ON-cells are inhibited. (B) Average firing rate for OFF-(left) and ON-cells (right) during presentation of different small chirps. There appears to be greater difference in averaged responses for OFF-cells than ON-cells. (C) Indeed OFF-cells can discriminate between varying small chirps on high frequency beats.

### Encoding of aggressive signals does not efficiently represent chirp properties

Small chirp stimuli cause abrupt phase-shifts in a low frequency beat cycle (Fig 1A). The spatial geometry of the electric field and electroreceptors is such that a stimulus that increases electrical amplitude on one side of the body will cause a decrease in amplitude on the other side, eliciting a response from ipsilateral ON-cells and OFF-cells respectively. As we have shown before (23), superficial ON-cells of the lateral segment of the ELL respond with a more salient response than other cells in the ELL: they produce a high-frequency stereotyped burst of spikes after the chirps that serves as a particularly effective chirp-detection mechanism. While both bursting and non-bursting responses contain some information about the amplitude modulation (AM) shape of the chirp stimulus there is still high error even when pooling the responses for 13 cells (Fig 1B). Importantly, this information about AM shape does not allow an observer to discriminate differences in the parameters of the emitted chirps such as duration or frequency rise (Fig 1C). For these stimuli, a chirp X with a specific duration and frequency occurring at a given phase will cause a different AM than the same chirp occurring at a different phase (compare stimuli in Fig 1A). Consequently, two chirps that differ in parameters could elicit responses that are more similar to each other than to the response of the same chirp occurring at different phases. A decoder cannot rely on spiking pattern to estimate variation in any variable parameter of these chirp signals as both duration and frequency information are obscured by the phase parameter which is not controlled by the emitting fish (24). In the context of natural behavior the signal’s characteristics hinder discrimination and as a result is encoded with a synchronized bursting code that is efficient for detection (23) but less so for discrimination (Fig 1C).

### Chirp properties are accurately encoded in high-frequency beat contexts

Big and small chirps occurring on high frequency beats span several cycles of the beat and thus the overall AM shape of the signals is virtually unaffected by the phase at which the chirp started (25). Big chirps cause a transient increase in beat frequency that is accompanied by a decrease in the beat amplitude directly proportional to the frequency increase (24). These chirps cause a graded increase in firing rate in OFF-cells and a cessation of firing in ON-cells (Fig 1D). The increase in OFF-cell firing can be of only a few Hz in some cells or of a few hundred Hz in others (21). For chirps that differ in duration ON and OFF cells, both allow accurate discrimination based on the responses of 12 cells (Fig 1E), the duration of the response being proportional to the duration of the chirp. When presented with two chirps that do not differ in duration but only in frequency, this pause in ON-cell firing cannot allow accurate discrimination. OFF-cells respond in a graded manner, increasing firing in relation to both chirp duration and frequency increase, allowing for efficient discrimination even when chirp duration is identical (Fig 1F). These data show that big chirps on high frequency beats are encoded with spiking patterns that allow the identification and discrimination of chirp characteristics.

Even though small chirps and big chirps are categorically different signals that often mediate different behaviors (19, 20), their behavioral impact (26–28) and the way they are encoded in the electroreceptors (29) depends on the beat frequency(25). Our data also suggest that the ELL responses to small chirps on high frequency beats are more similar to big-chirps than small chirps on low-frequency beats. OFF-cells respond with a variable increase in firing and ON-cells with a pause in firing (Fig 2A). The small chirps shown in Figure 2 cause an average increase in firing of 20 Hz in OFF-cells, which corresponds to a doubling of baseline frequency. This increase appears slightly stronger and longer in the larger of the small chirps (Fig 2B). Similar to the response of OFF-cells to big chirps (21), this population response is relatively invariant from trial to trial, therefore the difference in response shape between two chirps is larger than the typical variability in the response to a single given chirp. This is quantified by the discrimination error level, which reaches very low values for OFF-cell population responses of 12 neurons (Fig 2C). Also like the response to big chirps, ON-cell responses differ for the various small chirps but those differences are small relative to the trial-to-trial response variability leading to inaccurate discrimination coding. It is possible that larger populations (>>13 cells) could lead to error-less discrimination as the variability in the population response could be partially averaged out. Therefore, we conclude that both ON-cells and OFF-cells could support the discrimination of small chirps on high frequency beats but that OFF-cells do so more efficiently.

### Correlating behavior with coding strategy

Our neurophysiological data argue for a match between a signal with a structure that prevents chirp properties from being readily discriminated (i.e. small chirps occurring on low frequency beats) and a neural code that provides reliable detection, but does not support efficient discrimination: synchronous bursting (Fig 1B). In contrast, signals with structures better suited for discrimination are encoded with graded, heterogeneous responses that efficiently carry information about varying chirp characteristics. However, the neurophysiological data cannot demonstrate is that this change in coding strategy in fact mediates different perceptual tasks. In the following experiments, we test the hypothesis that behavioral discrimination can occur only in the context of a high frequency beats, Consistent with predictions made from the observed coding strategies, behavioral responses to small chirps on low frequency beats should be indiscriminant of chirp properties.

To test behavioral discrimination performance, we used a paradigm that has been used successfully in a wide variety of animals and modalities (30–33), a habituation-dishabituation assay. Animals respond most strongly to novel stimuli and habituate to a stimulus presented repeatedly. In the habituated state, the animal responds relatively weakly or not at all to the stimulus but when a novel stimulus is presented the animal displays a resurgence of response that reflects the fact the stimulus is perceived as novel. This dishabituation response -or lack thereof-demonstrates that the animal discriminated -or not-between the first and second stimulus. One of the strengths of this assay is that it takes advantage of the animal’s innate response to a stimulus rather that forcing an artificial behavior onto the animal (e.g. classical conditioning). The test therefore reveals how perceptual abilities are used to guide innate behaviors. In this case, sensory information about a stimulus’ novelty is used to drive an increase in behavioral response. Brown ghost knifefish placed in a confined space (here, an aquarium of 30 cm ×30 cm ×10 cm) will react to a conspecific signal by increasing swimming movement and speed, swimming around the source of the stimulus as if “investigating” and sometimes chirping, biting or lunging in that direction (Fig 3A). Our stimulation regime leads to habituation in the first 60 minutes. After 90 minutes of stimulation with a given chirp, the stimulus was switched (Fig 3B) to a stimulus with the same beat frequency and the same chirp type but different chirp parameters (frequency rise or duration) as used in the neurophysiological experiments.

**Figure 3:**
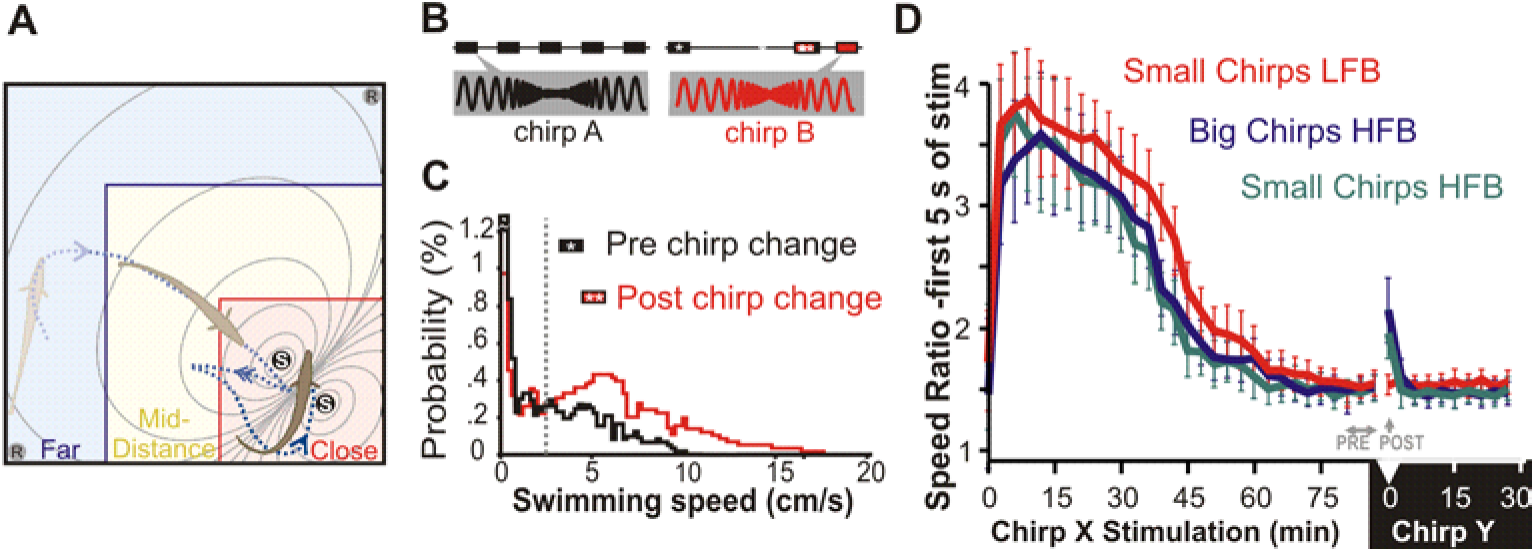
Chirps on high frequency beats cause dishabituation (A) Schematic of the experimental tank. Recording electrodes (R) recorded electrical activity. Fish were classed as being Far, Mid, or Close based on distance from stimulating electrodes (S). (B) Chirp stimuli were played 1 min on/2 min off for 90 min (black boxes). After 90 min, a chirp of the same type (small/big) but with different frequency and duration was played on the same beat (red boxes). Asterisks mark stimulation bouts used for analysis. (C) Example of swimming speed increases as the result of dishabituation. To normalize for baseline activity level, a threshold (dotted vertical line) was determined at which there were an equal number of points on either side (time moving slow : time spent moving fast) for each fish. (D) The first 5 seconds of behavior during chirp playback show speed increases for novel chirps on high frequency beats. MANOVA followed by Tukey HSD showed 0.02<p<0.04 between the high frequency beat stimuli at time 0 of chirp Y and the last three stimulus presentations of chirp X (n=80 to 85 trials each). For trials with low frequency beat stimuli (n=85), the same comparisons give 0.27<p<0.45.

### Swimming speed changes in response to chirps on high frequency beats

Various aspects of the behavioral response were quantified and several showed some habituation-dishabituation effect (biting, lunges, chirping; data not shown). Counting lunges or chirps clearly showed habituation but only revealed dishabituation in a small subset of individuals. If the dishabituation response is not strong enough to cross the threshold where it would produce one of these highly aggressive behaviors the measure would not reveal the phenomenon. We found swimming speed and changing distance from the stimulus source be more sensitive measures of dishabituation.

The distribution of swimming speeds during a given stimulation bout typically had a bimodal form with a peak at slower swimming speed indicative of a passive state and the faster swimming speed occurring during active swimming. We take the ratio of time spent in these two states (normalized for each fish; see SI for details) to quantify the strength of the behavioral response. The ratio will be higher than 1 at the beginning of the assay when the fish reacts most strongly (i.e. swims faster) to the stimulus and also if the fish exhibits dishabituation to the novel stimulus.

Swimming speed decreased markedly in the first hour of stimulation and there was no difference in average swimming speeds across stimulus type at the beginning of the stimulation protocol (ANOVA for time 0 of chirp X stimulation: p=0.3). The first stimulus bout with a new chirp led to a small but not significant increase in speed ratio for both small and big chirps on high frequency beats and no increase for small chirps on low frequency beats (Fig S2). Two factors affect the magnitude of this dishabituation effect: 1-The dishabituation response is marked in the first few seconds of stimulation but disappears quickly. Averaged across the whole minute of stimulation the short dishabituation effect does not influence values much; 2-dishabituation is observed more clearly in some fish than others as a function of the position of the fish when the new stimulus starts (see next paragraph and Fig 4A). To counter the short-lived response time we quantified speed ratios in the first 5 seconds of each bout (Fig 3D). These 5-second samples show a similar habituation trend that plateaus after 60 minutes. The dishabituation when the new stimulus is played is obvious (see figure legend for statistics) for chirps of both types on high frequency beats but not for small chirps on low frequency beats.

**Figure 4:**
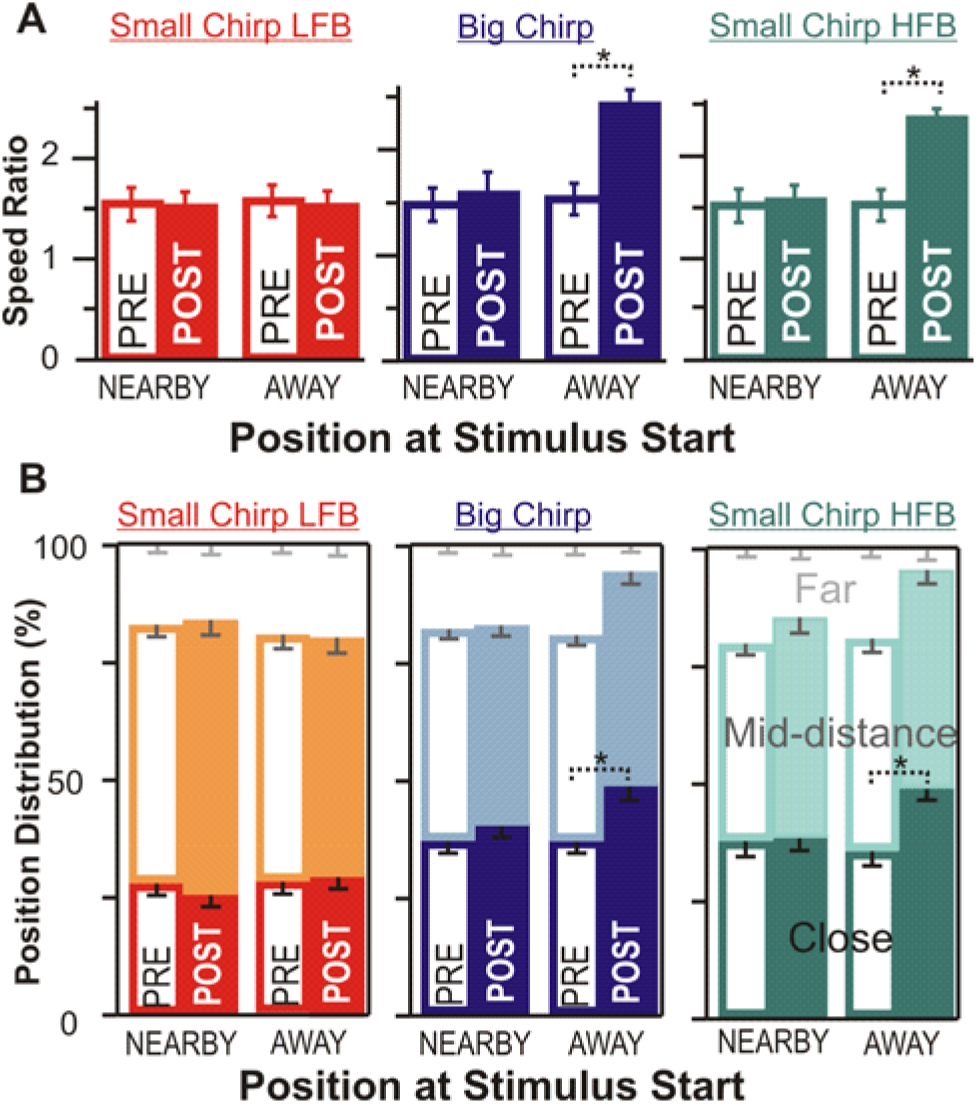
Speed change and position change are both dependent on location. (A) Fish close to the stimulating electrodes (NEARBY) show no change in swimming speed in response to any novel chirps. Fish farther from the stimulating electrodes at novel chirp start (AWAY) increase speed only for chirps on high frequency beats. Dotted brackets with asterisks indicate significant differences; paired t-test, p< 0.01 for the high frequency beat stimuli “away” and p>0.1 for all others. (B) Using the first five seconds after novel chirp stimulation, fish farther from stimulating electrodes at the start of the novel chirp moved closer to the stimulating electrodes for chirps on high frequency beats. Dotted brackets with asterisks indicate significant differences in the time spent close to the stimulus; paired t-test, p<0.01.

### Location affects dishabituation response

When identifying factors that influence dishabituation, we observed that some fish did not show a recovery of active swimming when the new stimulus was played. In every case, the fish was close to the stimulation electrode just prior to the start of the 1-minute playback. In contrast, fish that were positioned further from the electrode more reliably recovered active swimming when a new chirp on a high frequency beat was delivered. We quantified the position effect by separating the trials based on the position of the fish in the 1 second prior to the start of the new stimulus. Trials where the fish had over 50% of their body length in the “far” zone (Fig 3A) were categorized as being “away” from the stimulus, whereas fish in either remaining zone were defined as being “nearby”. We contrasted the speed ratio in the three stimulation bouts pre-stimulus-change (only the first 5 seconds of stimulation are used, as in Fig 3D) to the swimming speed in the first seconds of post-chirp-change stimulation. Our data confirm that dishabituation is observable only if the fish is not in close proximity to the stimulation electrode (Fig 4A, statistics in figure legend).

Position in the tank also reveals changes in behavioral responsiveness of the fish if the fish remains close to the stimulus source when actively engaged by the stimulus and less so when the behavioral response habituates. A measurement of distance in a small tank between a long fish and a stimulus source that is, not one, but two points is hard to define. To quantify position in a way that is not subject to the choice of two points defining stimulus and fish locations, we categorized the position based on zones (see Fig 3A, SI). Although we repeated our analysis with other ways of quantifying distance (e.g. nose to middle of dipole distance) with qualitatively similar results. The overall fish position distribution did not change as a result of habituation (Fig S3). When habituated the fish often remained still on one of the sides of the tank (near or far from the electrodes) leading to a broad distribution of positions. By again selecting the first 5 seconds of each stimulus and categorizing the trials based on position just prior to the novel stimulus we can see a change in the position of the fish for stimuli on high frequency beats (Fig 4B). This is consistent with the scenario that leads to the results in Figure 4A: in the first few seconds of a new high frequency beat stimulus, if the fish is far from the electrode it swims towards the electrode and passes over it several times presumably to investigate the source of the new stimulus, indicating that it has perceived the chirps as two distinct stimuli.

## DISCUSSION

An ideal coding system balances reliable detection of a signal with minimal error and effective discrimination between closely related stimuli to perform efficiently during behavioral tasks (34). However, balancing these goals may drive a tradeoff between efficient detection of a signal and detailed discrimination between two similar signals (13, 35, 36). In *A. leptorhynchus* we argue that two separate coding strategies developed as a way to maximize both efficient categorical perception as well as discrimination of detailed information about chirp structure dependent on which need is most behaviorally relevant in a given context.

### Neural coding is matched to context dependent signal structure

Small chirps occurring on low frequency beats cause large shifts in the background beat, but vary dramatically in the shape of the electrical image dependent on which phase of the beat they occur on. This phase variance masks properties of small chirps such as duration and frequency rise. Emitting fish do not actively control which phase of the beat they chirp on (24). Likewise, a given small chirp presented at different phases is encoded in an invariant manner in the midbrain and elicits invariant behavioral responses in the receiving fish (37). The lack of phase control in chirp behavior, and invariance of response in the physiology demonstrates that phase is most likely not relevant to either the emitting or receiving fish and actively hinders the ability to discriminate chirps.

The transient disruptions caused by a small chirp on the slow ongoing background frequency cause a brief synchronization in the firing or quiescence of the electroreceptor afferents (29, 38). Likewise, ON-cells respond with a stereotyped burst of action potentials when presented with small chirps at the beat trough, and OFF-cells respond when the chirp occurs at the beat peak (23). The brief durations of small chirps in conjunction with the long cycles of slow frequency beats hinder discrimination based on chirp duration and frequency rise. Our data confirm the results of Marsat and Maler (21) and Marsat et al. (23) demonstrating that responses to small chirps on low frequency beats cannot be discriminated from one another by a biologically realistic decoding mechanism (39).

At higher beat frequencies both big and small chirps span more than one beat cycle, reducing the effect of phase on the electrical image the fish receives. This change in the signal structure may allow the animal to encode details about chirp duration and frequency more accurately. At these beats electroreceptor afferents phase lock more synchronously, and all chirps disrupt that synchrony (25, 29). Desynchronization in afferent firing may give rise to the observed ON-cell inhibition and variable increase in OFF-cell firing that occurs in response to both chirp types on high frequency beats. This graded OFF-cell response provides sufficient information for an ideal observer to accurately discriminate between big chirps (21). Here, we demonstrate that this coding strategy is not specific to big chirps, but is also incited by small chirps. The change in chirp structure and as a result the change in coding is a product of the social context (beat frequency) rather than due to the properties of the chirp itself.

### Perceptual ability corresponds to coding strategy

The structure of a communication signal limits coding, but the behavioral relevance of the signal may also play a role in how a signal is processed. We argue that the type of neural codes used in each situation should be matched to the structure of the signal and the perceptual task being performed. Changing behavioral states, noise present in an environment, and social context may all influence the needs of the organism, and thus how a signal is best encoded. In diverse sensory systems including vision, audition, and touch there are demonstrable links between the coding of a stimulus and the animal’s sensory acuity(7, 40–42). Our behavioral results show not only the differential coding of chirps but systemically correlates that code to different behavioral outcomes. Only chirps presented on high frequency beats produced a dishabituation response indicative of chirp discrimination, while small chirps on low frequency beats induced no behavioral changes. We cannot exclude the possibility that fish perceived a difference in small chirps on low frequency beats but did not react; however, it does not affect the strength of our conclusion. The fact that the behavioral response is determined by the presence or absence of the chirp -not by its detailed characteristics-demonstrates that the behavior is guided by chirp detection. Whether the information to discriminate is present at one point in the nervous system is irrelevant, as this information does not influence behavior.

These results closely match the observed neural coding patterns. The match between signal structure, neural code and behavioral performance could be interpreted as one of these aspects (e.g. signal structure) causing the other to be shaped accordingly. Instead, we put forth that our results argue for the specialization of the sensory system for specific perceptual tasks, either detection or discrimination.

### Specialization of coding for separate tasks

In *A. leptorhynchus* beat frequency establishes the social context of conspecific interaction and may function as a filter, priming the ELL for efficient, task-dependent signal processing. Different sub-populations of cells best encode the signals to be detected versus the signals to be discriminated (ON-cells vs. OFF-cells respectively; (21, 23)). However, these studies and our current results also demonstrate that both subpopulations’ responses allow detection and discrimination. Therefore, the difference in processing is not mainly a difference in the subpopulation that carries out the task, but rather a difference in response pattern and stimulus shape. In other words, the neural coding strategy changes as a function of the stimulus. Ollerenshaw et al. (13) demonstrated in the rodent vibrissae system that adaptation changes the neural code from one primed for detection to one better able to discriminate, a change mirrored in behavioral performance. In this case, changes in coding are not implemented by adaptation but are a consequence of the fixed properties of the network such as feedforward frequency tuning (25) and feedback (16). As indicated by our results and studies of coding strategies in diverse sensory systems (Visual (43, 44); insect auditory (45, 46)), adaptive changes in coding specialized for detection and discrimination tasks (47) might be a wide-spread phenomenon that allows efficient coding in a constantly changing environment.

Burst firing is a particularly robust method of encoding sensory information (14). This feature detection scheme is especially useful when encoding transient, but important communication signals (23) or small prey items (48). In this light, the stereotyped burst response produced by small chirps on low frequency beats may reflect their ethological significance to the animal. On low frequency beats, indicative of same sex interactions between closely sized animals, small chirps are most commonly produced during agonistic encounters, and are often used for establishing territorial dominance (20, 22). Behavioral studies indicate that chirp timing is a key determinant of the interaction they mediate (49). Presumably, the chirp emission pattern rather than the detailed structure of each chirp carries the information to guide behavioral responses. Detailed discrimination of these chirps may be sacrificed for reliability in detection. The need for an efficient detection code might not be obvious in a laboratory setting when two fish engage in close-range aggression but this task is artificially easy compared to natural interactions. Root masses and other obstructions, distance and eavesdropping scenarios, the presence of multiple conspecifics, movement of various nearby signals, healthiness of the sensory surface (the skin) and various other factors can make the chirp-detection task harder. In these settings it may be beneficial to have a readily detectable signal that indicates conspecific presence efficiently, rather than the more intense bouts of courtship chirping.

Burst structure is stereotyped and in large part dictated by the burst mechanism. While some stimulus features may be extracted from bursts (50) they are limited in the amount of information they can encode in their structure due to that stereotyped structure. Discrimination tasks therefore benefit from an alternate form of coding. High entropy population responses with graded heterogeneous responses are well suited for that purpose since they accomplish signal whitening (21) and thus maximize channel capacity (51, 52). The heterogeneous responses of the population may sacrifice the reliability of burst coding for the detailed coding of chirp features. As beats greater than 100 Hz are generally indicative of male-female interactions, chirps produced in this context are more likely to be courtship-related than agonistic (19, 22). For such interactions, it is possible that the need to assess mate fitness or establish individual identity becomes more valuable than reliable detection. Switching from a burst code to a graded one may enhance the ability to differentiate between subtly different chirps.

## METHODS

See SI for full methods

### Neurophysiology

All methods were approved by WVU IAUCUC (Protocol 1512000009). Surgical methods are based on those described in Marsat, Proville, and Maler (23) and Marsat and Maler (21). Fish of both sexes were used. *In vivo* recordings were made via extracellular electrodes (Frank & Becker, 1964). Pyramidal cells of the lateral segment of the ELL were targeted and identified by location in relation to major surface blood vessels, depth from the surface of the brain, as well as neural response properties (53, 54).

### Stimulation during neurophysiology

All stimuli were sampled at 20 kHz and created using MATLAB. Stimulus strength was adjusted to provide ~20% contrast as measured close to the fish’s skin.

Chirp stimuli consisted of a Gaussian frequency modulation of the background beat presented once per second on either a 10 or 120 Hz beat. The frequency and duration of chirps were chosen to mimic a range of natural signals (55). Details about chirp properties can be found in SI methods.

### Neural data analysis

Analysis of discrimination is based on Marsat and Maler (21) and modifications to methods originally described by van Rossum (56). This method quantifies how similar or dissimilar spiking patterns are using firing rate and spike timing. Jackknife resampling was used to calculate the standard errors displayed in Figures 1 and 2.

### Behavioral paradigm

Recordings were acquired from 250 trials using fish of both sexes. Stimuli were played via a pair of submerged electrodes located in one quadrant of the tank (Fig 4A) at a strength replicating the average fish’s electric field. Fish were played one of three chirp stimuli matched those used in the neurophysiology experiment. These chirps were played at a rate of twice per second for one minute, followed by a two-minute break. This stimulation pattern was repeated for 90 minutes. After 90 minutes a chirp of the same category (either small or big) but with a different frequency increase and duration was played on the same beat frequency (Fig 3B).

### Behavioral Analysis

The video files were analyzed in MATLAB. For each 1-minute stimulus bout, the distribution of swimming speeds was calculated (Fig 3C) based on the distance moved between each video frames and the resulting data were used as described in the Results section.

## Acknowledgments

Funding was provided by West Virginia University and the National Science foundation grant IOS-1557846. The authors would like to thank Dr. Len Maler for his assistance and resources for a pilot study related to this study and Drs Andrew Dacks and G. Troy Smith for helpful comments on the manuscript.

## SUPPLEMENTAL INFORMATION

### METHODS

#### Neurophysiology

Surgical methods are based on those described in Marsat, Proville, and Maler (1) and Marsat and Maler (2). Fish of both sexes were anaesthetized and respirated with a solution of Tricaine-S (Tricaine Methanesulfonate, Western Chemical, Inc.) in water (.25g/L) for the duration of the surgery. After the application of a local anesthetic (Lidocaine HCl 2%, Hospira, Inc.) skin and overlying soft tissues were removed from a small area of the skull. A portion of the exposed skull was glued to a fixed post for stability while the portion of the skull overlying the ELL was then removed. Fish were immobilized with a . 1mL injection of Tubocurarine Chloride Pentahydrate (.2mg/mL, TCI), switched to anesthetic free water for respiration, and allowed to recover from surgery for approximately 20 minutes in the experimental tank before stimulation and recording. The experimental tank (40 × 45 × 20 cm) contained water matched to the home system (26-27° C, 250-300μS). *In vivo* recordings were made via metal filled extracellular electrodes (Frank & Becker, 1964) and amplified (A-M Systems, Model 1700), and data recorded (Axon Digidata 1500 and Axoscope software) at a 20kHz sampling rate. Pyramidal cells of the lateral segment of the ELL were targeted and identified by location in relation to major surface blood vessels, depth from the surface of the brain, as well as neural response properties (3, 4).

#### Stimulation during neurophysiology

All stimuli were sampled at 20 kHz and created using MATLAB (Mathworks, Inc.). Pyramidal cells respond to changes in EOD amplitude (4, 5), so stimulation was provided by a direct modulation of a carrier frequency matching the fish’s own rather than by mimicking a second EOD. This method allows for tight control over the signal the fish receives and is commonly used in similar experiments (6, 7). The baseline EOD was recorded via electrodes near the head and tail of the fish. Each EOD cycle triggered a sine wave generator (Rigol DG1022A) to generate one cycle of a sine wave matched to the animal’s own. This signal was then multiplied using a custom-built signal multiplier (courtesy of the Fortune Laboratory, New Jersey Institute of Technology) by the AM stimulus to create the desired modulation of the electric field around the fish. It was played through a stimulus isolator (A-M Systems, Model 2200) into the experimental tank via two 30.5cm electrodes arranged parallel to the fish’s longitudinal axis. This arrangement stimulates the majority of electroreceptors equally. The stimulus strength was adjusted to provide ~20% contrast (the difference between the maximum and minimum of amplitude modulation divided by the baseline EOD).

Chirp stimuli consisted of a Gaussian shaped frequency modulation of the background beat presented once per second on either a 10 or 120 Hz beat. The frequency and duration of chirps were chosen to mimic a range of natural signals (8). Small chirps were either 10ms long with a 60 Hz increase, or 15ms long with a 122 Hz increase. Big chirps were either 15 ms long with a 300 Hz increase, or 45 ms long with a 900 Hz increase. Chirp frequency increase was tied to a signal amplitude decrease of 0.08% for each Hz of frequency rise, based on natural chirp properties (9). All four chirps were played on both the 10 and 120 Hz beats but responses to big chirps on the 120 Hz beat were not examined in detail. Small chirps were presented at several different phases of the beat, typically at the peak or trough of the sine wave.

#### Neural data analysis

Analysis of discrimination is based on Marsat and Maler (2) and modifications to methods originally described by van Rossum (10). This method accounts for both the firing rate as well as the temporal pattern of spikes to quantify how similar or dissimilar spiking patterns are. Our analysis is mathematically equivalent to previous approaches (11–14) but instead of displaying a confusion matrix and calculating the amount of information carried by the population response about stimulus identity we display the ROC curves and quantify an error level.

Spike trains were binarized and convolved with an *α* filter, *f* (*t*) = *t*^−2.45/*τ*^, with τ being the width of the function at half maximum (15). A portion of the result *R*(*t*) was extracted for analysis, specifically, a window around the timing of the presented chirps (–15 to 30 ms relative to the middle of big chirps and –10 to 30 ms relative to small chirps) as shown in figure 1A. The distance *D_xy_* between the two spike trains *x* and *y* of length *L* is defined as: 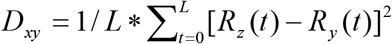 (Fig S1 B). Larger distances indicate more dissimilar spike trains. In addition to the response of individual neurons, we looked at population responses by averaging several spike trains using the function 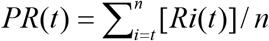. The result (*PR*(*t*)) represents a population of neurons presented with the same stimulus and mimics a neuron integrating postsynaptic potentials with similar weights (16). These population responses were created by randomly pairing multiple individual responses, simulating the response of a diverse population of cells. Up to 200 random combinations of spike trains from all recorded neurons were used for all comparisons. Responses to different chirps (X vs. Y) were compared as well as multiple responses to the same chirp (X vs. X). Distance (*D_XY_* or *D_xx_*) was calculated for all sets of combined responses, *PR*_x_(*t*) and *PR*_y_(*t*), creating an array of response distance for each comparison. The probability distributions of the values in these arrays (*P*(*D*_*XY*_) or *P*(*D*_*xx*_)) were used for ideal observer analysis (Fig S1 C). Receiver operating characteristic curves were generated by varying the threshold distance for discrimination, T. For each threshold value, the probability of non-discrimination (*P*_D_) is calculated as the sum of *P*(*D*_XY_ > *T*), and the probability of false discrimination (*P*_F_) as the sum of *P*(*D*_XX_ > *T* (Fig S1 D). The error level for each threshold value is *E* = 1/2*P*_F_ + 1/2(1 – *P*_D_). The error in discrimination reported in the figures are the minimum values of *E* (Fig 1F). This measure of error rate is closely related to the Kullback-Leibler divergence quantifying how much overlap two distributions have. These measures thus quantify how different two distributions are without relying on assumptions of normality as other statistical tests often do. Jackknife resampling (leaving one neuron at a time out of the analysis) was used to calculate the standard errors displayed in Figures 2–4.

#### Behavioral paradigm

The experimental tank measured 30 cm x 30 cm x 10 cm and contained water matched to the fish’s home tank (26-27°C and a conductivity of 200-300 μS). Fish were allowed to acclimate overnight. The experimental tank was shielded from light and illuminated with an infrared light source and filmed with an infrared camera (Logitech C920, from which its IR filter was removed). Video was captured at 24 frames/s and a spatial resolution of 1280×720 pixels (the tank covered the width of the frame). Stimuli were played via a pair of submerged electrodes (dipole spacing=10 cm) located in one quadrant of the tank (Fig 3A) at a strength replicating the average fish’s electric field. Fish were played one of three chirp stimuli: big chirps on a 120 Hz beat, small chirps on a 10 Hz beat, or small chirps on 120 Hz beat. Chirp properties were as described for the neurophysiology experiments. These chirps were played at a rate of twice per second for one minute, followed by a two-minute break with no stimulation. This stimulation pattern was repeated for 90 minutes to habituate the fish to the chirp. After 90 minutes a chirp of the same category (either small or big) but with a different frequency increase and duration was played on the same beat frequency (Fig 3B). Behavior was assessed throughout the experiment when the stimulus was on to be able to quantify the habituation and possible dishabituation of the response.

#### Behavioral Analysis

Recordings were acquired from 250 trials. A given fish was tested only once a day and could be tested up to three times total: once with each stimulus. The video files were imported in MATLAB where a custom program was used to analyze the video frame-by-frame. The semi-automatic analysis identified the position of three points on the fish: the tip of the nose, the tip of the tail and the 2D center of mass.

These points were determined automatically by the program but visual inspection of the results was required to correct occasional errors (e.g. flipping of the tail and nose). Swimming speed and position could easily be calculated given the frame time stamp and the spatial calibration. The results displayed took into account the position of the nose, but the center point gave similar results; results using the tip of the tail were not evaluated. For each 1-minute stimulus bout, the distribution of swimming speeds was calculated (Fig 3C) based on the distance moved between each video frames and the resulting data were used as described in the Results section.

---

**SI1.**
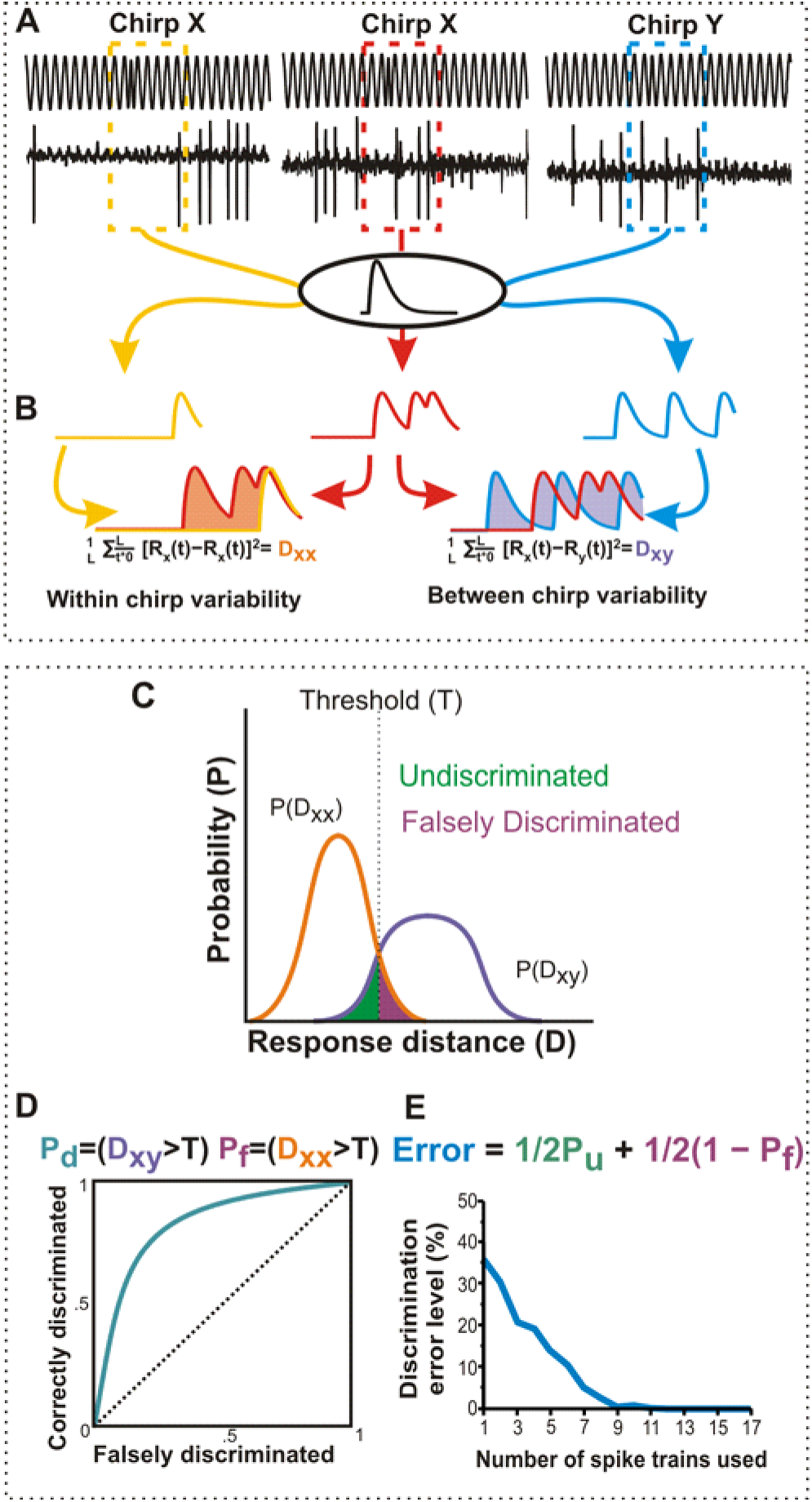
Discrimination analysis. (A) Representative chirp response spike trains. A portion of the recording before and after chirp presentation is extracted for analysis. Spike trains are binarized, then convolved with an α filter. This process can be performed on individual spike trains, or averaged spike trains to simulate a population response. (B) The response distance is the difference between multiple filtered responses. Responses to the same chirp determine firing variability (D_xx_) while responses to different chirps are analyzed to determine between chirp response distance (D_xy_). (C) These individual or population response distances are then used to generate response probability distributions. Overlap between these two distributions represents error in discriminating between two different chirps: either the probability of going undiscriminated (a false negative, P_u_) or the probability of being falsely discriminated (false positive, P_f_). (D) From this we generate receiver operating characteristic curves to determine the performance of an ideal observer attempting to distinguish chirps from the responses. The threshold for discrimination (T) can be varied to achieve the best performance to maximize the number of chirps correctly discriminated (P_d_) while minimizing the number falsely discriminated. (E) Once the best threshold is determined, errors in discrimination are calculated and displayed as a percentage of incorrectly sorted responses out of all responses. This process is repeated for increasing numbers of neurons in our population averages to determine how population heterogeneity improves performance.

**S2:**
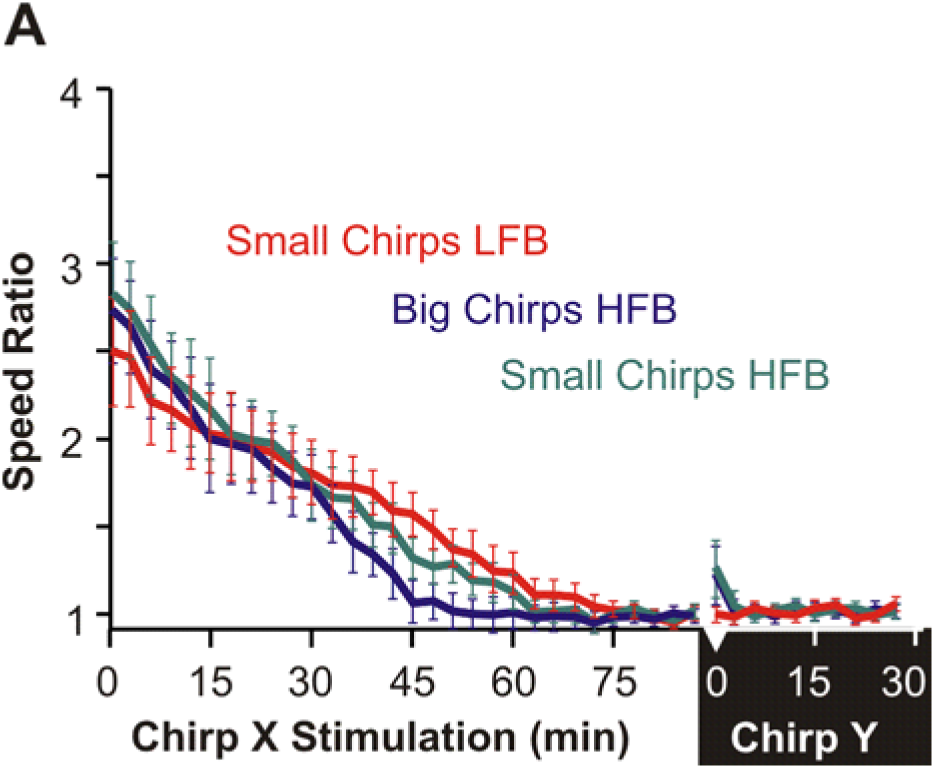
(A) Plot of speed ratio (fast/slow swimming) through the duration of the experiment averaged across trials for each 60s chirp playback. MANOVA followed by Tukey HSD showed 0.15<p<0.32 between the HFB stimuli at time 0 of chirp Y and the last three stimulus presentations of chirp X. For LFB stimuli, the same comparisons give 0.36<p<0.42.

**S3:**
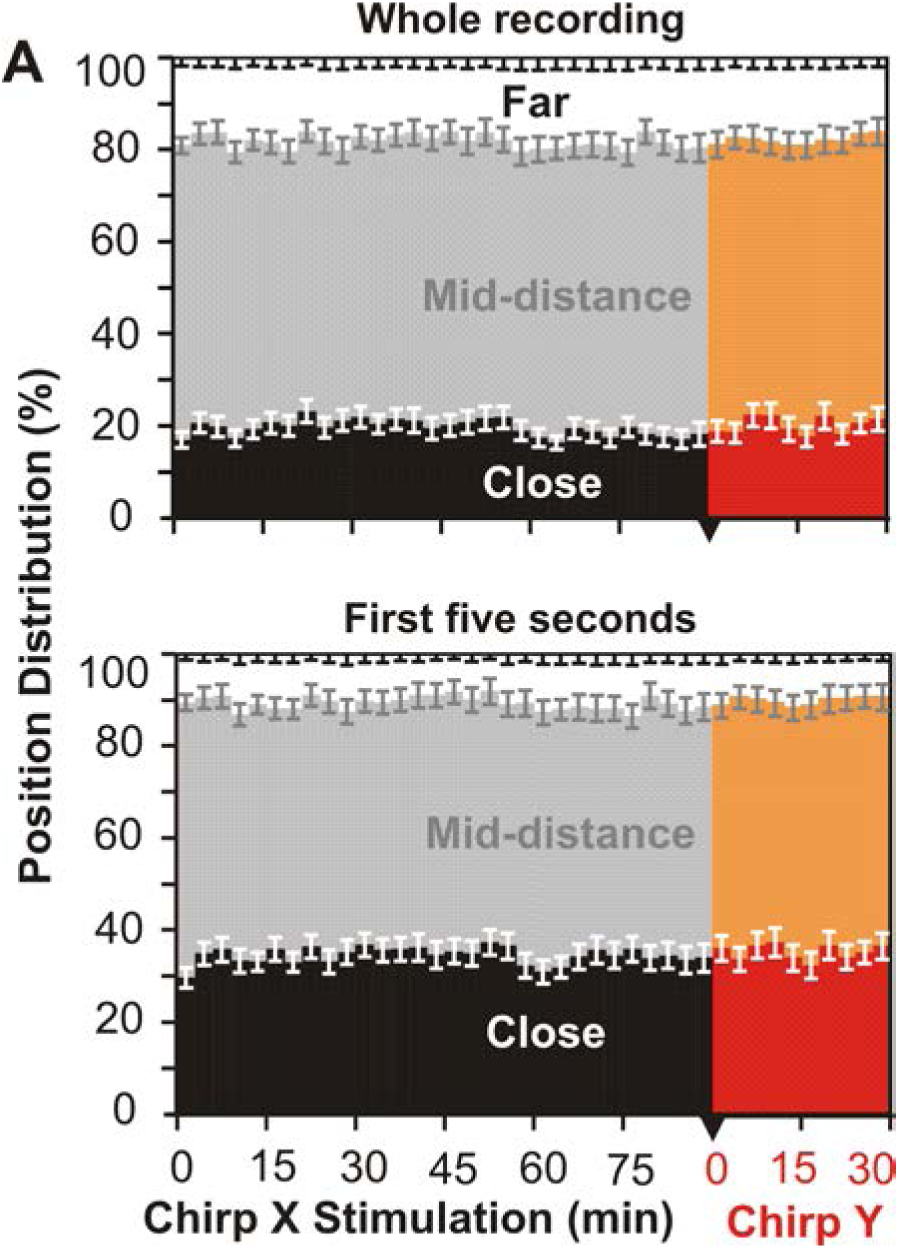
Position in tank does not change with habituation. (A) The population maintained distribution throughout the course of the experiment for both small chirps on low frequency beats (shown) and both chirps on high frequency beats (data not shown). (B) Overall distribution did not change after playing the novel chirp stimulus, even when using only the first five seconds of the recording.

## REFERENCES

1. Richards DG (1981) Alerting and Message Components in Songs of Rufous-Sided Towhees. Behaviour 76(3/4):223–249.

2. Bradbury JW, Vehrencamp SL (2011) Principles of animal communication (Sunderland: Sinauer Associates). Second.

3. Kröger BJ, Birkholz P, Kannampuzha J, Neuschaefer-Rube C (2011) Categorical perception of consonants and vowels: Evidence from a neurophonetic model of speech production and perception. Lect Notes Comput Sci (including Subser Lect Notes Artif Intell Lect Notes Bioinformatics) 6456 LNCS:354–361.

4. Mitchell BR, Makagon MM, Jaeger MM, Barrett RH (2006) Information content of coyote barks and howls. Bioacoustics 15(3):289–314.

5. Oglesbee E, Kewley-Port D (2009) Estimating vowel formant discrimination thresholds using a single-interval classification task. J Acoust Soc Am 125(4):2323–35.

6. Bradbury, J.W. and SLV (2011) Principles of animal Communication.

7. Arabzadeh E, Petersen RS, Diamond ME (2003) Encoding of Whisker Vibration by Rat Barrel Cortex Neurons: Implications for Texture Discrimination. Jounal Neurosci 23(27):9146–9154.

8. Freedman DJ, Riesenhuber M, Poggio T, Miller EK (2001) Categorical representation of visual stimuli in the primate prefrontal cortex. Science (80-) 291(5502):312–6.

9. Tremblay F, Ageranioti-Belanger SA, Chapman CE (1996) Cortical mechanisms underlying tactile discrimination in the monkey. I. Role of primary somatosensory cortex in passive texture discrimination. J Neurophysiol 76(5):3382–3403.

10. von Heimendahl M, Itskov PM, Arabzadeh E, Diamond ME (2007) Neuronal activity in rat barrel cortex underlying texture discrimination. PLoS Biol 5(11):e305.

11. Fairhall AL, Lewen GD, Bialek W, de Ruyter Van Steveninck RR (2001) Efficiency and ambiguity in an adaptive neural code. Nature 412(6849):787–92.

12. Moore CI (2004) Frequency-dependent processing in the vibrissa sensory system. J Neurophysiol 91(6):2390–9.

13. Ollerenshaw DR, Zheng HJ V, Millard DC, Wang Q, Stanley GB (2014) The adaptive trade-off between detection and discrimination in cortical representations and behavior. Neuron 81(5): 1152–64.

14. Krahe R, Gabbiani F (2004) Burst firing in sensory systems. Nat Rev Neurosci 5(1): 13–23.

15. Marsat G, Pollack GS (2012) Bursting neurons and ultrasound avoidance in crickets. Front Neurosci 6(JULY):1–9.

16. Marsat G, Longtin A, Maler L (2012) Cellular and circuit properties supporting different sensory coding strategies in electric fish and other systems. Curr Opin Neurobiol 22(4):686–692.

17. Panzeri S, Macke JH, Gross J, Kayser C (2015) Neural population coding: combining insights from microscopic and mass signals. Trends Cogn Sci 19(3): 162–72.

18. Tripathy SJ, Padmanabhan K, Gerkin RC, Urban NN (2013) Intermediate intrinsic diversity enhances neural population coding. Proc Natl Acad Sci U S A 110(20):8248–8253.

19. Hagedorn M, Heiligenberg W (1985) Court and spark: electric signals in the courtship and mating of gymnotoid fish. Anim Behav 33:254–265.

20. Hupé GJ, Lewis JE (2008) Electrocommunication signals in free swimming brown ghost knifefish, Apteronotus leptorhynchus. J Exp Biol.

21. Marsat G, Maler L (2010) Neural heterogeneity and efficient population codes for communication signals. J Neurophysiol 104(5):2543–55.

22. Henninger J (2015) Social interactions in natural populations of weakly electric fish.

23. Marsat G, Proville RD, Maler L (2009) Transient signals trigger synchronous bursts in an identified population of neurons. J Neurophysiol 102(2):714–23.

24. Zupanc GKH, Maler L (1993) Evoked chirping in the weakly electric fish Apteronotus leptorhynchus : a quantitative biophysical analysis. Can J Zool 71(11):2301–2310.

25. Walz H, Grewe J, Benda J (2014) Static frequency tuning accounts for changes in neural synchrony evoked by transient communication signals. J Neurophysiol 112(4):752–65.

26. Dunlap KD, Larkins-Ford J (2003) Production of aggressive electrocommunication signals to progressively realistic social stimuli in male Apteronotus leptorhynchus. Ethology 109(3):243–258.

27. Triefenbach FA, Zakon HH (2008) Changes in signalling during agonistic interactions between male weakly electric knifefish, Apteronotus leptorhynchus. Anim Behav 75(4):1263–1272.

28. Triefenbach F, Zakon H (2003) Effects of sex, sensitivity and status on cue recognition in the weakly electric fish Apteronotus leptorhynchus. Anim Behav 65(1):19–28.

29. Benda J, Longtin A, Maler L (2006) A synchronization-desynchronization code for natural communication signals. Neuron 52(2):347–58.

30. Cheney DL, Seyfarth RM (1988) Assessment of meaning and the detection of unreliable signals by vervet monkeys. Anim Behav 36(2):477–486.

31. Miller-Sims VC, Bottjer SW (2012) Development of Auditory-Vocal Perceptual Skills in Songbirds. PLoS One 7(l2).

32. Penn D, Potts WK (1998) Untrained mice discriminate MHC-determined odors. Physiol Behav 64(3):235–243.

33. Zuberbühler K, Cheney DL, Seyfarth RM (1999) Conceptual semantics in a Nonhuman Primate. J Comp Psychol 113(1):33–42.

34. Barlow HB (1961) Possible principles underlying the transformations of sensory messages. Sensory Communication, ed Rosenblith WA (MIT Press, Cambridge, MA), pp 217–234.

35. Guido W, Lu SM, Vaughan JW, Godwin DW, Sherman SM (1995) Receiver operating characteristic (ROC) analysis of neurons in the cat’s lateral geniculate nucleus during tonic and burst response mode. Vis Neurosci 12(4):723–741.

36. Reinagel P, Godwin D, Sherman SM, Koch C (1999) Encoding of visual information by LGN bursts. J Neurophysiol 81(5):2558–2569.

37. Metzen MG, Hofmann V, Chacron MJ (2016) Neural correlations enable invariant coding and perception of natural stimuli in weakly electric fish. Elife 5:1–22.

38. Benda J, Longtin A, Maler L (2005) Spike-frequency adaptation separates transient communication signals from background oscillations. J Neurosci 25(9):2312–21.

39. Larson E, Billimoria CP, Sen K (2009) A biologically plausible computational model for auditory object recognition. J Neurophysiol 101(1):323–31.

40. Adibi M, Arabzadeh E (2011) A comparison of neuronal and behavioral detection and discrimination performances in rat whisker system. J Neurophysiol 105(1):356–65.

41. Bendor D, Wang X (2007) Differential neural coding of acoustic flutter within primate auditory cortex. Nat Neurosci 10(6):763–71.

42. von Heimendahl M, Itskov PM, Arabzadeh E, Diamond ME (2007) Neuronal activity in rat barrel cortex underlying texture discrimination. PLoS Biol 5(11):e305.

43. Lesica, Nicholas A., Stanley GB (2004) Encoding of Natural Scene Movies by Tonic and Burst Spikes in the Lateral Geniculate Nucleus. J Neurosci 24(47):10731–10740.

44. Sherman SM (2001) Tonic and burst firing: dual modes of thalamocortical relay. Trends Neurosci 24(2):122–126.

45. Marsat G, Pollack GS (2004) Differential temporal coding of rhythmically diverse acoustic signals by a single interneuron. J Neurophysiol 92(2):939–48.

46. Sabourin P, Pollack GS (2010) Temporal coding by populations of auditory receptor neurons. J Neurophysiol 103(3):1614–21.

47. Stanley GB (2013) Reading and writing the neural code. Nat Neurosci 16(3):259–263.

48. Oswald A-MM, Chacron MJ, Doiron B, Bastian J, Maler L (2004) Parallel processing of sensory input by bursts and isolated spikes. J Neurosci 24(18):4351–62.

49. Hupé GJ (2012) Electrocommunication in a Species of Weakly Electric Fish Apteronotus Leptorhynchus: Signal Patterning and Behaviour. Dissertation (Université d’Ottawa/University of Ottawa).

50. Marsat G, Pollack GS (2010) The structure and size of sensory bursts encode stimulus information but only size affects behavior. J Comp Physiol A Neuroethol Sensory, Neural, Behav Physiol 196(4):315–320.

51. Doi E, Lewicki MS (2014) A Simple Model of Optimal Population Coding for Sensory Systems. PLoS Comput Biol 10(8).

52. Simoncelli EP, Olshausen BA (2001) Natural image statistics and neural representation. Annu Rev Neurosci 24(1): 1193–1216.

53. Maler L, Sas E, Johnston S, Ellis W (1991) An atlas of the brain of the electric fish Apteronotus leptorhynchus. J Chem Neuroanat 4(1):1–38.

54. Saunders J, Bastian J (1984) The physiology and morphology of two types of electrosensory neurons in the weakly electric fish Apteronotus leptorhynchus. J Comp Physiol A 154(2):199–209.

55. Bastian J, Schniederjan S, Nguyenkim J (2001) Arginine Vasotocin Modulates a Sexually Dimorphic Communication Behavior in the Weakly Electric fish Apteronotus leptorhynchus. J Exp Biol 204(11):1909–1923.

56. van Rossum MC (2001) A novel spike distance. Neural Comput 13(4):751–763.

## REFERENCES

1. Marsat G, Proville RD, Maler L (2009) Transient signals trigger synchronous bursts in an identified population of neurons. J Neurophysiol 102(2):714–23.

2. Marsat G, Maler L (2010) Neural heterogeneity and efficient population codes for communication signals. J Neurophysiol 104(5):2543–55.

3. Maler L, Sas E, Johnston S, Ellis W (1991) An atlas of the brain of the electric fish Apteronotus leptorhynchus. J Chem Neuroanat 4(1):1–38.

4. Saunders J, Bastian J (1984) The physiology and morphology of two types of electrosensory neurons in the weakly electric fish Apteronotus leptorhynchus. J Comp Physiol A 154(2):199–209.

5. Bastian J (1981) II. The effects of moving objects and other electrical stimuli on the activities of two categories of posterior lateral line lobe cells in Apteronotus albifrons. J Comput Physiol 144:481–494.

6. Bastian J, Heiligenberg W (1980) Neural correlates of the jamming avoidance response of Eigenmannia. J Comp Physiol 136(2):135–152.

7. Benda J, Longtin A, Maler L (2005) Spike-frequency adaptation separates transient communication signals from background oscillations. J Neurosci 25(9):2312–21.

8. Bastian J, Schniederjan S, Nguyenkim J (2001) Arginine Vasotocin Modulates a Sexually Dimorphic Communication Behavior in the Weakly Electric fish Apteronotus leptorhynchus. J Exp Biol 204(11):1909–1923.

9. Zupanc GKH, Maler L (1993) Evoked chirping in the weakly electric fish Apteronotus leptorhynchus : a quantitative biophysical analysis. Can J Zool 71(11):2301–2310.

10. van Rossum MC (2001) A novel spike distance. Neural Comput 13(4):751–763.

11. Laubach M, Wessberg J, Nicolelis MA (2000) Cortical ensemble activity increasingly predicts behaviour outcomes during learning of a motor task. Nature 405(6786):567–71.

12. Stecker GC, Harrington IA, Middlebrooks JC (2005) Location coding by opponent neural populations in the auditory cortex. PLoS Biol 3(3):e78.

13. Tremere LA, Pinaud R (2011) Brain-generated estradiol drives long-term optimization of auditory coding to enhance the discrimination of communication signals. J Neurosci 31(9):3271–89.

14. Vonderschen K, Chacron MJ (2011) Sparse and dense coding of natural stimuli by distinct midbrain neuron subpopulations in weakly electric fish. J Neurophysiol 106(6):3102–18.

15. Machens CK, et al. (2003) Single auditory neurons rapidly discriminate conspecific communication signals. Nat Neurosci 6(4):341–342.

16. Larson E, Billimoria CP, Sen K (2009) A biologically plausible computational model for auditory object recognition. J Neurophysiol 101(1):323–31.

